# Ontology-guided segmentation and object identification for developmental mouse lung immunofluorescent images

**DOI:** 10.1101/2020.06.26.174367

**Authors:** Anna Maria Masci, Scott White, Ben Neely, Maryanne Ardini-Polaske, Carol B. Hill, Ravi S Misra, Bruce Aronow, Nathan Gaddis, Susan E. Wert, Scott M. Palmer, Cliburn Chan, LungMAP Consortium

**Author notes:** (Equal contribution).

## Abstract

**Background:** Immunofluorescent confocal microscopy uses labeled antibodies as probes against specific macromolecules to discriminate between multiple cell types. For images of the developmental mouse lung, these cells are themselves organized into densely packed higher-level anatomical structures. These types of images can be challenging to segment automatically for several reasons, including the relevance of biomedical context, dependence on the specific set of probes used, prohibitive cost of generating labeled training data, as well as the complexity and dense packing of anatomical structures in the image. The use of an application ontology surmounts these challenges by combining image data with its metadata to provide a meaningful biological context, and hence constraining and simplifying the process of segmentation and object identification.

**Results:** We propose an innovative approach for the automated analysis of complex and densely packed anatomical structures from immunofluorescent images that utilizes an application ontology to provide a simplified context for image segmentation and object identification. We describe how the logical organization of biological facts in the form of an ontology can provide useful constraints that enhance automatic processing of complex images. We demonstrate the results of ontology-guided segmentation and object identification in mouse developmental lung images from the Bioinformatics REsource ATlas for the Healthy lung (BREATH) database of the Molecular Atlas of Lung Development (LungMAP1) program.

**Conclusion:** The microscopy analysis pipeline library (micap) is available at https://github.com/duke-lungmap-team/microscopy-analysis-pipeline. Code to reproduce our analysis of LungMAP images is also available at https://github.com/duke-lungmap-team/lungmap-pipeline. Finally, the application ontology is available at https://github.com/duke-lungmap-team/lung_ontology and includes example SPARQL queries.

**Contact:** Anna Maria Masci

## Background

We describe the development of ontology-guided segmentation and object identification for immunofluorescent images from the Molecular Atlas of Lung Development Program (LungMAP) consortium^1^. LungMAP seeks to advance research in lung development and disease by creating, integrating and disseminating genomic and imaging data sets. The immunofluorescent confocal microscopy data set includes hundreds of diverse images spanning multiple stages of development and stained with distinct probe combinations. At an early embryonic stage, the developing lung parenchyma is a densely packed tissue, whereas the mature lung exhibits an open, filamentous alveolar structure. Some structures are present only during specific developmental stages, while other structures undergo drastic anatomical and physiological changes over time. The combination of antibody probes in a subset of images is chosen to target particular cells within an anatomical structure to study these changes and their relevance to the developing lung. Consequently, images acquired at different developmental stages and with different probe combinations display significant visual differences. The limited number of images sharing the same visual appearance for a given anatomical structure poses significant challenges to their segmentation and identification, particularly in the prohibitive cost of generating labeled training data. We use an application ontology to incorporate the metadata representing the varations in image acquisition to provide a meaningful biological context, making the process of segmentation and object identification more manageable.

To identify specific structures in immunofluorescent microscopy images of the developing lung, a human observer would make use of experimental, imaging and anatomical contextual information. The experimental context includes information such as species and developmental stage. The imaging information includes the magnification and labeled probes used. Finally, the anatomical context includes the list of structures typically found in the appropriate developing stage of the lung that are identifiable using a particular set of probes, as well as the hierarchical nesting of substructures within structures and spatial relationships between structures. Our application ontology captures the types of contextual data a human observer would use, in a way that can be incorporated into the image processing workflow.

An ontology is a controlled, structured vocabulary that provides logical definitions of terms representing distinct categories and their relationships in a particular domain. With strict control of unique term identifiers and logical consistency, ontologies enable the interoperability of heterogeneous data and are widely used as a tool to facilitate the sharing of knowledge. For example, the Gene Ontology^2, 3^ (http://geneontology.org/) has been used to annotate large collections of literature and data, facilitating data integration and discovery. In turn, this has led to the development of software that can query diverse sets of experimental data curated by different groups of experts and stored in independent databases. In the biomedical disciplines, ontologies provide rigorous, unambiguous descriptions of biological entities and the relationships among them using standardized and well understood formats (http://ontology.buffalo.edu/biomedical.htm). The Open Biomedical Ontology (OBO) Foundry (http://obofoundry.org) ontologies are designed to describe orthogonal biological features, utilizing a common standard to ensure interoperability. The OBO Foundry consists of foundational ontologies that define the terms of interest in a domain. Application ontologies combine and take advantage of the terms defined in the foundational ontologies for use in a specific application.

Several methods have been proposed for segmentation of fluorescence microscopy images based on computer vision algorithms^4-12^. We propose the creation of an application ontology to provide biomedical context in a machine-readable way to increase the power of computer vision algorithms for the identification of complex objects within microscopy images. While ontologies have been previously utilized for providing automatic image annotation^13^, our novel approach utilizes an ontology to dynamically create segmentation parameters targeting structures likely to be found in a particular image. Our approach combines image metadata, including the probes used, with a newly developed ontology to dynamically define constraints allowing the targeted segmentation of candidate regions. These candidate regions are then processed to calculate a series of feature metrics used to classify the regions into distinct anatomical structures, which can be further segmented to identify individual cells within each structure. The ontology defines the possible structures identifiable at each developmental stage and provides precise labels allowing for interoperability with other ontologies.

An alternative to classical computer vision is the use of deep learning with convolutional neural networks (CNN)^14-17^. Examples of well-known CNNs include Fast R-CNN^18^ and Mask R-CNN^19^. For biomedical image segmentation, U-NET^20^ is a popular architecture. However, U-NET only provides semantic segmentation into connected pixels, whereas our application calls for instance segmentation into distinct objects. To prevent overfitting, these methods require a large number of labeled training images, which are difficult to achieve in many practical contexts. For example, CNNs for image classification and object detection often use training databases with thousands or even millions of images. Several strategies exist to address the lack of adequate training data in microscopy experiments, including attempts to restore lower quality images for inclusion in training data^21^ and data augmentation. The immunofluorescent images produced by the LungMAP consortium are generated as part of specific experiments and hence heterogeneous. For a given probe set (i.e., a fluorescent antibody, or antibodies, used to recognize a specific molecule(s) associated with a distinct cell type), only a small number of images are available, and we encountered no images that required excluding due to image quality issues. When evaluating various neural networks, we attempted various data augmentation techniques as well as transfer learning using the COCO dataset^22^, but were unable to coax standard CNN models, such as Mask R-CNN, to perform competitively with careful feature engineering using classical image analysis methods. Even if more images were available, the burden of constructing a large training set would be prohibitive since the images require experts in developmental lung anatomy for accurate labeling.

## Results

### Image Selection

The immunofluorescent images within the LungMAP project are heterogeneous, including variation in species, developmental age, magnification and labeled probe combination used. We first used the ontology categories to group images into image sets with the same species, development age, magnification, and probe combination. This process is automated using the LungMAP SPARQL API, which returns this grouping metadata in the Resource Description Framework (RDF) labeled graph format for each queried image. The grouping simplifies image segmentation, since images within a set are now homogeneous with respect to extrinsic variables.

### Creation of training data

Manual segmentation of training data is time-consuming and prone to subjective contour boundaries. Most manual segmentation tools also require the user to provide the region labels, potentially requiring review and modification of the labels to match the those in the classification pipeline. To address these issues, we developed a Graphical User Interface (GUI) utility to create training data (Figure 6). Since the pipeline’s segmentation and classification processes are independent, the segmentation routine, including image pre-processing, can be used to create regions. Additionally, the ontology allows us to filter the list of possible anatomical structures for a given development age and the combination of probes present in the image set.

**Figure 1:**
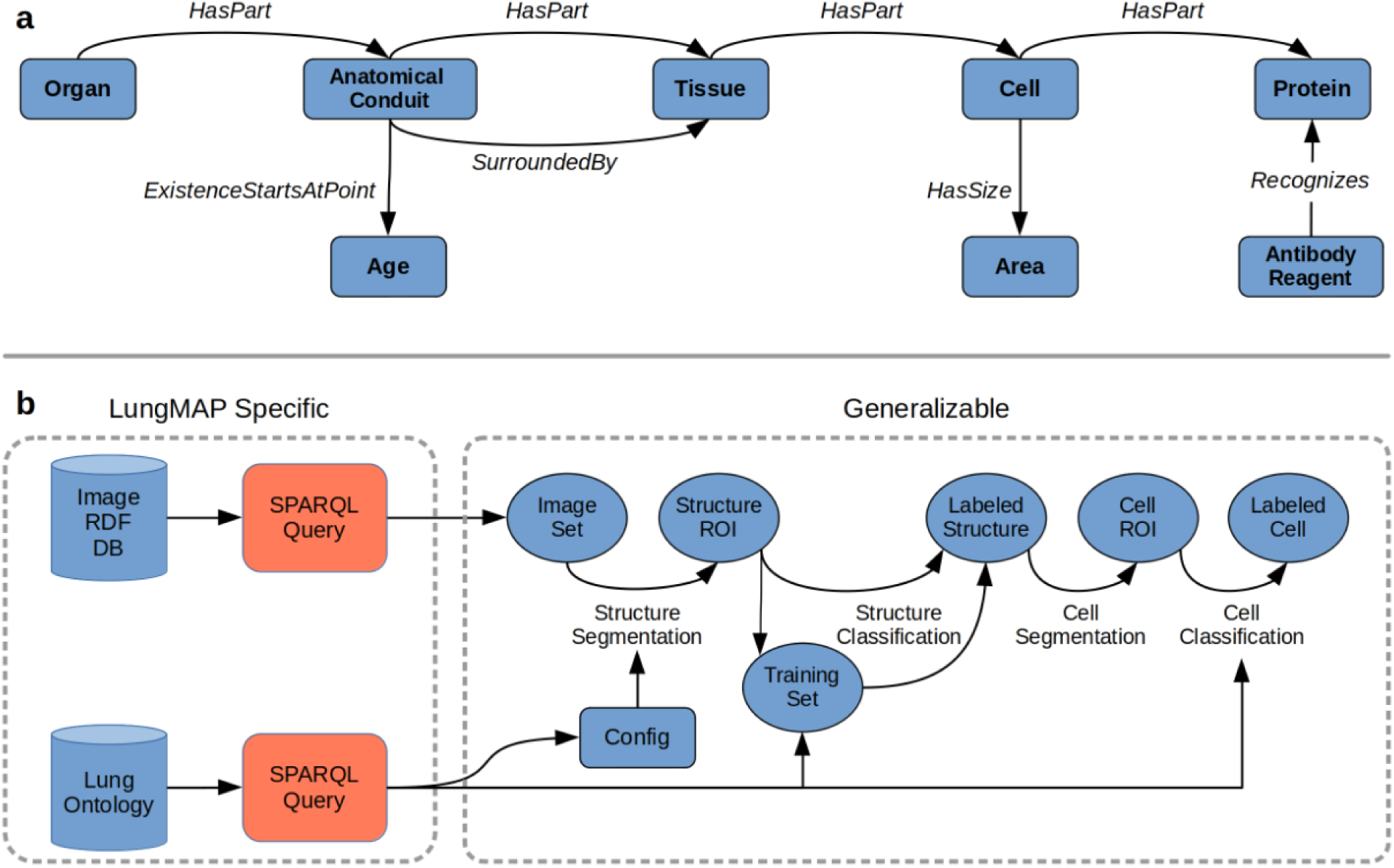
a) Structure of application ontology for determining the relationships between antibodies and anatomical entities (cells, tissues, etc.) in fluorescence microscopy images. b) Integration of ontology queries and application ontology into workflow of image segmentation and classification.

**Figure 2:**
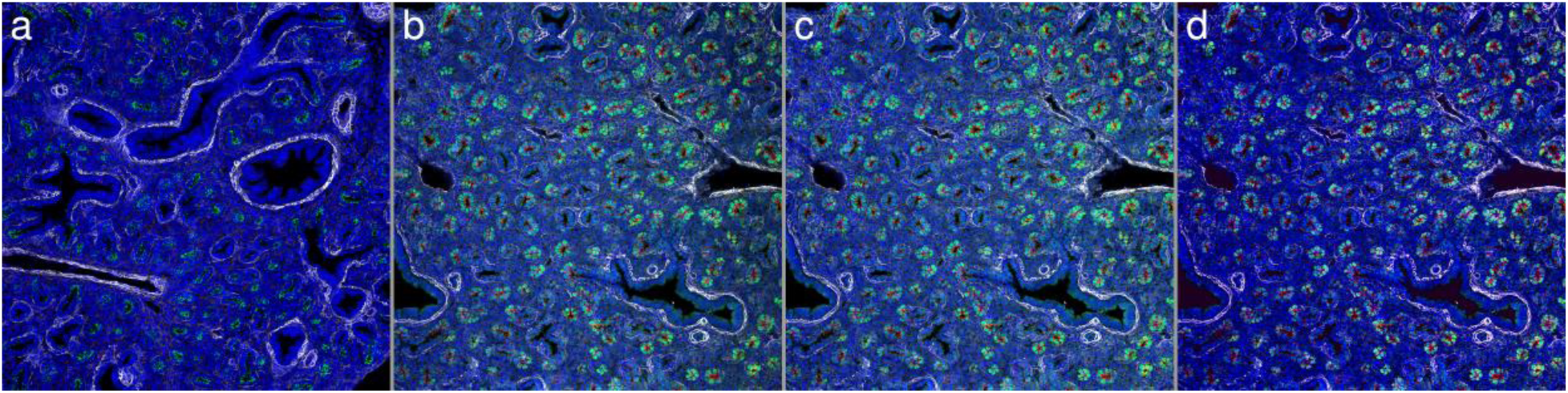
Image pre-processing. a) Reference image chosen from an image set for color correction. b) Original appearance from a non-reference image in the image set. c) Non-reference image after gradient correction. d) Non-reference image after color correction (and gradient correction).

**Figure 3:**
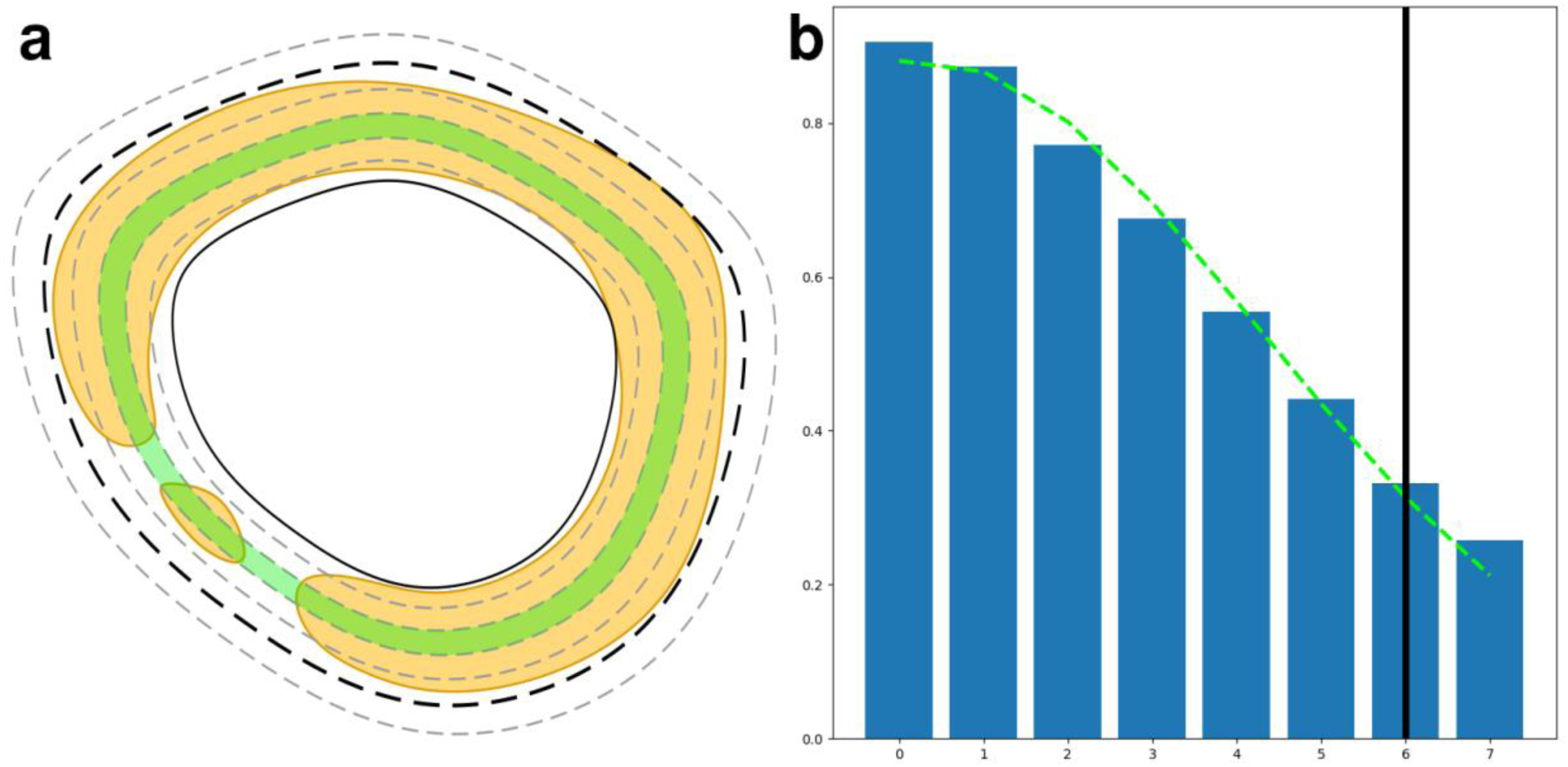
a) Illustration of dilating a candidate region using the signal mask. The yellow region represents the signal mask, an area likely to contain an anatomical structure. The signal mask for a single anatomical entity is often disconnected, illustrating the infeasibility of using it directly. The solid black line outlines the original candidate boundary. The dashed grey lines represent successive dilations. The green ring represents the dilation iteration with the maximum signal percentage. Signal values from each ring are fitted with a Gaussian and the final boundary (black, dashed line) is determined at 1.5 standard deviations. b) Plot of signal percentages from successive dilations of a real candidate region. The green dashed line is the Gaussian fit. The solid black line marks the iteration for the final contour size.

**Figure 4:**
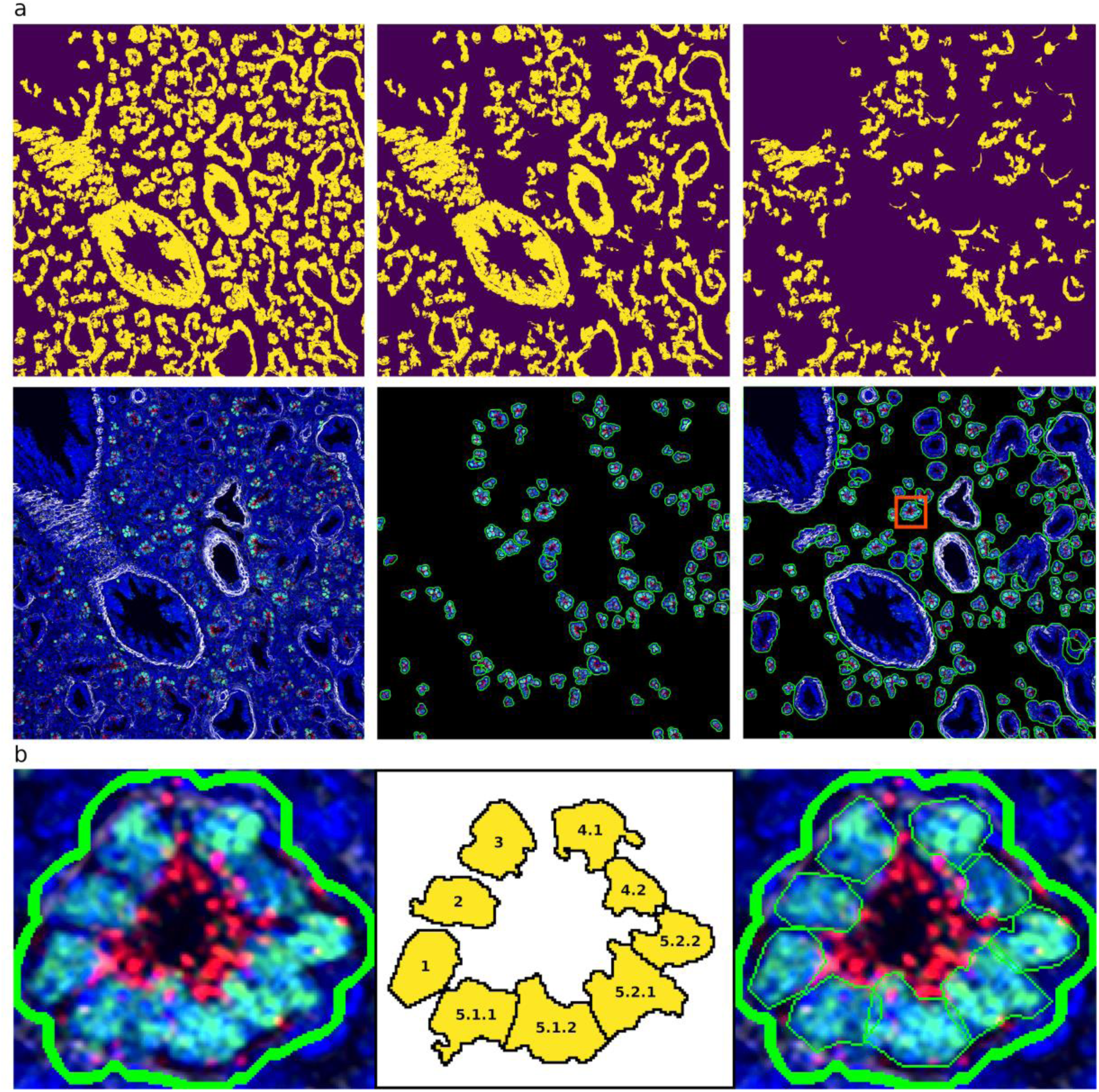
a) Stages in automated structure segmentation. First row shows signal mask progression: top-middle after color segmentation stage, top-right is the residual signal mask after all stages. Second row: left shows original image, middle image shows candidate regions found in color segmentation stage, right shows all final candidates. The red box highlights a single region to demonstrate cell segmentation b) Segmentation of cells within structure with recursive spectral cluttering. From left to right: original structure from structure segmentation algorithm, order of segmentation of cells into structures in 3 recursive stages, cell segments within structure segment.

**Figure 5:**
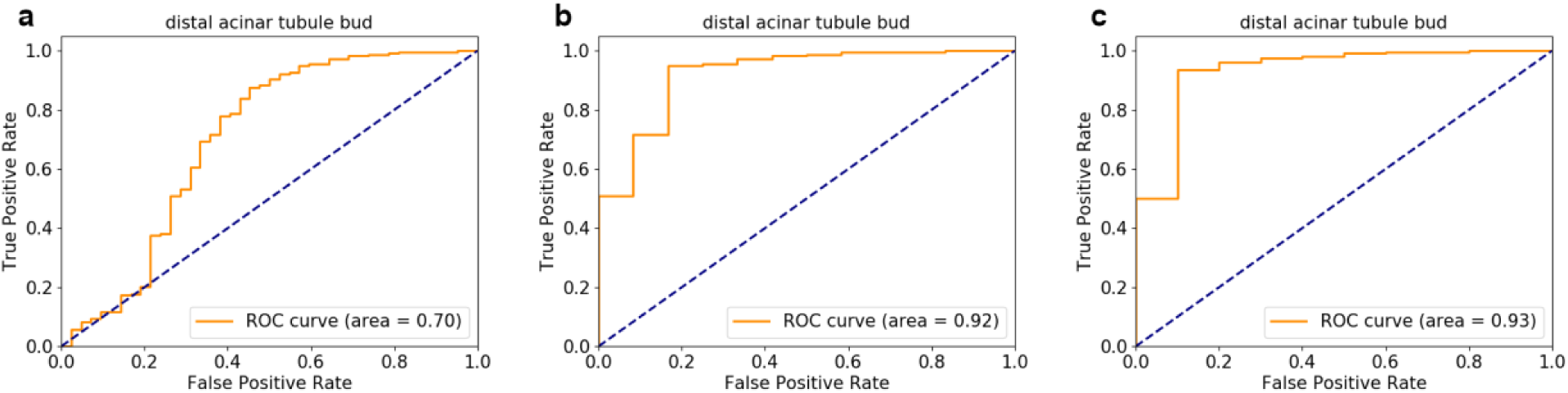
ROC curves for labels generated by XGBoost as compared with ground truth labels assigned by a histopathologist. The probe set in the selected images targeted the distal acinar tubule bud. The three curves show results based on different acceptance criteria for whether a region correctly identifies the structure. A) Only a single region with >50% overlap of a whole structure as labeled by the pathologist is accepted as correct. All other sub-regions, regardless of label, are marked incorrect. B) When the histopathologist-labeled structure includes more than one region, these regions are manually merged, and the single merged region is accepted as correct. C) When the histopathologist-labeled structure includes more than one region, the individual regions are all accepted as correct.

**Figure 6:**
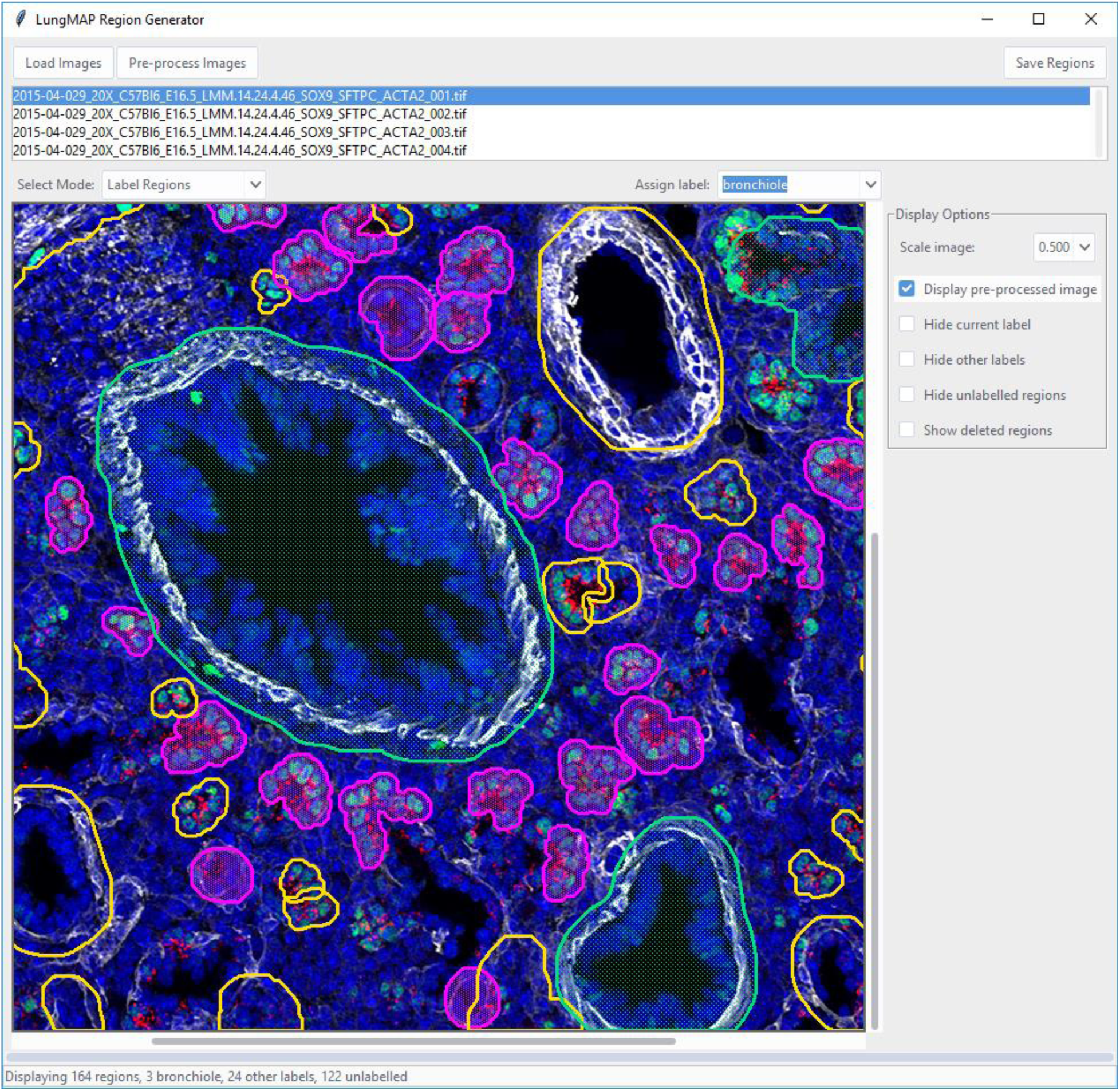
Screenshot of application for labeling segmented structures which integrates the application ontology and image analysis pipeline. The application has multiple modes accessible via drop-down box, including modes for labeling, deleting, splitting, and new regions.

The tool provides an interface for an expert to visualize the segmented structure candidates generated and accept or reject them. Labels from an ontology-constrained label set can be easily applied to the accepted regions. Regions covering more than one structure can automatically be split using a recursive binary segmentation routine. Finally, any structure missed by the automatic segmentation can be manually drawn as polygons, allowing images to be comprehensively segmented. This is an important feature as our training process is designed to automatically create a “background” class from non-segmented regions in these comprehensively segmented images.

### Targeted segmentation of structures

For a given image set, we use a SPARQL Protocol and RDF Query Language (SPARQL) traversal of the probe-protein-cell-tissue-structure graph of the application ontology to limit the classes of structures that are identifiable given a specific probe combination. In the LungMAP data sets, a particular set of probes (usually 3) is used to target a particular anatomical structure. The ontology allows ranking anatomical structures by the number of probes used in the image set that can label a candidate structure. The anatomical structures targeted by *the greatest number of* probes are prioritized as the primary object(s) to identify. In addition, the pipeline attempts to identify any other structures that have at least one targeting probe. For example, in an image of the lung from a mouse at the developmental stage E16.5 with the probe set ACTA2, SOX-9, and SFTPC, the target structure is the distal acinar tubule bud with 2 matching probes (SOX-9 and SFTPC), and the remaining identifiable anatomical structure classes with a single probe are the bronchiole, the proximal acinar tubule, the pulmonary artery, and the pulmonary vein. The application ontology also plays an important role in providing contextual information regarding the location of probes within structures, indicating whether a probe color is on the periphery of a structure or found within it (Supplemental Table 2).

The customizable color profile and the configurable segmentation stages provide flexible strategies for the analysis of a large range of immunofluorescent microscopic data sets. However, we found certain strategies more effective for the LungMAP application, and this drove the development of the ontology to automate the creation of segmentation configurations. For example, the first stage should target the most prominent visual structures found in the image set. These structures correspond to those that are targeted by the greatest number of probes. Querying the ontology provides this information.

We also found that knowing whether a probe targets the periphery, or the interior of a structure could further improve the segmentation results, and this information was then added to the ontology. Structures targeted by a probe that is found in the periphery of a structure are best placed at the end of the segmentation stage sequence. The rationale for this is that if placed at an earlier position of the segmentation sequence, then the more visually subtle structures in the image could be partially segmented. Using saturation-based segmentation for the intermediate stages proved more effective at completely segmenting these more concealed structures, with the priority towards larger structures. Larger structures are targeted with more aggressive (larger) blur kernels.

To label the candidate ROIs, a labeled training set is necessary, and the use of classical machine learning requires only a modest number of training examples for each structure class. The ROIs in the training data are labeled, the feature matrix is calculated for each ROI, and the resulting feature metric data is used to train an XG-Boost^23^ classifier. The trained classifier is then used to assign labels to new candidate regions. To evaluate accuracy, an image set of four images was selected to build four separate models in a leave-one-out approach. Next, using the held-out image and a trained model, we derived candidate regions and probabilities for each anatomical structure identified from the ontology. These candidate regions and their predicted anatomical structures were paired with the “ground truth” labeled by a histopathologist with expertise in developmental lung anatomy. These results are presented as ROC curves shown in Figure 5.

## Discussion

An application ontology can improve image segmentation and labeling by providing contextual information that is implicit when a human expert evaluates the image. We use the application ontology to construct flexible semantic queries using the SPARQL protocol to generate relevant contextual information for the image analysis pipeline. For immunofluorescent images of the developing lung, the context provided includes general information about the different developmental stages, the hierarchical organization of anatomical lung structures down to the cellular and molecular level, image metadata such as the magnification level, and the experiment-specific information of which fluorescent dyes are mapped to specific antibody probes. This contextual information constrains possibilities and simplifies inference. An additional benefit is that the application ontology guarantees that all terms are unique and derived from more foundational ontologies. This means that all labeled entities provide their own definitions and can be related to other entities. Using a standard vocabulary ensures consistent naming of structures following well-defined standards and also ensures that our labels can be unambiguously linked to other data sets and reference databases. For example, single cells identified in an image can be linked with other data for that cell type (e.g. scRNA-seq) in the BREATH databases, providing spatial resolution to other assay data. The ontology was developed collaboratively by an ontologist and a software developer, using an iterative process to ensure that each addition or modification to the ontology was driven by the intended use of the image analysis pipeline and could be queried programmatically.

There are limitations to using an ontology-driven approach for image segmentation and object classification. First, the construction of an application ontology is an iterative process and requires someone with expertise in biomedical ontologies collaborating closely with bioinformaticians who know how to use tools such as SPARQL for querying the ontology. Second, application ontologies are, by nature, constructed for a particular purpose, unlike more general domain (foundational) ontologies. Hence, elements of the application ontology must be adapted for new applications, for example, by adding new antibody specificities for a different probe set, or new anatomical relations for different organ systems. While the particular image processing routines described are specific to the lung development context, the framework of logical relationships established by this application ontology is generalizable to object identification in other biomedical imaging experiments. By providing context information in a machine-accessible format, application ontologies increase the feasibility of automatic image segmentation and object identification. Ontologies are commonly used in a static way for linkage and knowledge sharing, but image segmentation and object identification open new opportunities for ontologies to be used in a dynamic way for machine inference.

## Conclusion

In summary, application ontologies can be powerful tools to pair with immunofluorescent image processing, segmentation and object identification. In addition to their traditional virtues of providing a standard vocabulary and term definitions, logical inference over ontologies can make use of biomedical contextual information to constrain and simplify machine learning. We have demonstrated this approach for developmental lung studies, but believe that it is widely applicable, for example, to study cancer-immune interactions in the tumor microenvironment.

## Data Sharing

The microscopy analysis pipeline library (micap) is available on Github (https://github.com/duke-lungmap-team/microscopy-analysis-pipeline), and includes a basic example of usage using a trivial shapes data set. Code to reproduce our analysis of LungMAP images is also available on Github (https://github.com/duke-lungmap-team/lungmap-pipeline). Finally, the application ontology is available on Github (https://github.com/duke-lungmap-team/lung_ontology) and includes example SPARQL queries via the Python library “ontospy”.

## Methods

### Development of the ontology

The application ontology was developed following the Basic Formal Ontology (BFO, https://basic-formal-ontology.org/) formalism, and conforms to the requirements of OBO Foundry ontologies. The ontology was specified in the Web Ontology Language and developed using the Protégé (version 4.5) ontology editor. We searched OntoBee (http://www.ontobee.org/) and BioPortal (http://bioportal.bioontology.org/) for equivalent concepts in existing ontologies. To avoid duplication, we imported and used these equivalent terms. If no equivalent term was found, we created new terms and definitions and submitted them to an appropriate OBO Foundry ontology to consider for inclusion. We relied on Uberon^24^ for anatomical structures, Cell Ontology^25-27^ for cell types, Protein Ontology^28^ for proteins, Information Artifact Ontology^29^ for experimental protocol metadata, and the Relation Ontology^30^ for relations between entities. In addition, we ensured that every anatomical term in the application ontology mapped to a corresponding term in the LungMAP ontology^31^, which provides a comprehensive anatomical ontology of the developing lung.

Queries to the ontology are made using SPARQL. The application ontology contains the set of anatomical structures related to a specific antibody probe, including descriptions of the intermediate tissue and cell types that make up those structures. The ontology also contains common variations, or aliases, of the various probes commonly used in immunofluorescent microscopy and provides a standard unambiguous label for each probe.

All the relevant data elements were collected from the Bioinformatics REsource ATlas for the Healthy lung (BREATH) database and transformed into terms and relational expressions, mostly reused from other ontologies (Supplemental Table 1). The most critical metadata about the images are the fluorescent-labeled probes, which are antibodies against specific biological proteins. For this reason, we used a general class of Antibody Reagent from the Reagent Ontology (REO, http://www.ontobee.org/ontology/REO), and connected it to the targeted protein by the use of the relation *recognizes* adopted from REO and AntiO^32^. We then linked a specific cell type to the protein (recognized by the antibody reagent) via a *has_part* relation. For example, cell X *has_part* some protein Y and Antibody Reagent Z *recognizes* some protein Y. In turn, tissues link to cells, and higher-order anatomical structures link to tissues via the same relationship. The graph illustrating these relationships is shown in Figure 1a.

To capture the temporal nature of the development process, the ontology also includes age information. We used an *existence_starts_at_point* relation to capture the relationship between visible entities and age. This relation links an anatomical structure with the specific age at which that structure is developed and is visible within an image. As the images are from samples at only a few specific ages (e.g. E16.5), we used discrete categories for age. For other applications where a continuous time model is desirable, we would define overlapping intervals at which each entity can be found in the development process.

The ontology is used in the process of labeling training data, image segmentation, and mapping of image segments to named anatomical entities. When labeling training data, the ontology provides the list of possible anatomical structures for a given development age. For segmentation, the ontology provides contextual information regarding the location of probes, such as whether a probe is located on the periphery of a structure or found within it. For classification, the ontology constrains the anatomical entity candidate labels allowed given the inferred knowledge about the image and specific segment.

The inference over the ontology requires chained SPARQL queries between a set of probes and their related structures that can be time intensive. Hence, relationships between probes and structures are first determined using a set of SPARQL queries and then saved as a look-up table (LUT) for quicker access in the image analysis application. These LUTs can be re-generated programmatically when the ontology is updated. The integration of the ontology into the image processing pipeline is illustrated in Figure 1b.

### Development of image analysis pipeline

The image analysis pipeline we developed is tailored to the unique properties of immunofluorescent microscopy images, particularly targeting cases where regions of interest are densely packed and numerous. We chose to target these types of images as they are difficult to segment accurately using current computer vision pipelines and highly laborious to segment manually. The pipeline was developed in the Python programming language and is available as an open-source package named micap (**mic**roscopy **a**nalysis **p**ipeline).

Our pipeline makes use of metadata associated with each image sample, including species, development age, image magnification, names of the antibody probes, and the colors associated with those probes. Together, this information defines distinct image sets that are analyzed in separate runs of the pipeline. Training data for a single entity consists of an ontology-defined label and a list of vertices given as x-y coordinates that defines the boundary of a polygonal region.

### Development of feature metrics

Classical machine learning approaches require the reduction of pixel-level data to image feature vectors. The limited colors and gradients in immunofluorescent images can be usefully exploited to generate a limited set of quantitative feature metrics useful for distinguishing between biologically relevant segmentation classes. These features are based on color composition and distribution, region and sub-region shape statistics such as eccentricity and convexity, and simple texture statistics like hole area ratio (HAR).

We began the development of features using manually segmented regions and then extracted these regions into groups based on their labels. The extracted regions were visually inspected and analyzed using histograms of their HSV channels. The colors present in all of the images in the LungMAP database are blue: representing the nuclei stained with DAPI, black: representing non-cellular empty space, or “background” due to dark-field microscopy, and red, green, and white: typically used as fluorescent colors in immunofluorescent microscopy. Additionally, we observed that for small areas where a targeted antibody was expressed, the fluorescent color would interact with the blue DAPI stain to produce composite colors.

Based on these observations and the histograms of the hue channel in the regions we manually segmented, we partitioned the hue range into 3 main sections we term “major” colors, targeting red, green, and blue. We also binned the value channel into 3 monochromatic colors of black, gray, and white. Finally, we defined “minor” colors as the composite colors of violet, cyan, and yellow. With these 9 color categories established, the entire HSV space was partitioned such that any combination of hue, saturation, and value belongs to a unique color class. The complete definition of HSV ranges for the color classes are shown in Supplementary Table 3.

In total, there are 179 feature metrics calculated for each region, consisting of 19 metrics repeated for each of the 9 color classes and 8 global contour features. While the LungMAP data sets employ 3 fluorescence colors, more colors can be used in immunofluorescent microscopy^33^. With this in mind, we designed the analysis pipeline library to calculate feature metrics dynamically based on a configurable set of color definitions. By default, the library will use the 9 color definitions described above.

### Image pre-processing

Once an image set is selected for analysis, its images are preprocessed to be homogeneous with regard to intrinsic variables, such as illumination and color. In microscopy images, illumination inhomogeneity effects occur both within an image and between images. Intra-image variation in light intensity results from a focal light source and manifests as a non-uniform intensity across the field of view (FOV), with higher intensities near the center of the image and trailing out toward the edges. Intra-image FOV variation can be significant, with up to 30% variation in intensity from the center to the darkest edge^34^. Inter-image variation can occur in each of the hue, saturation and value channels. Differences in the placement and brightness of the focal light source, variations in the staining process, and photobleaching are among the sources contributing to intra-image variations of color and intensity. These effects are responsible for images that appear “washed-out” (low saturation values) or images where the DAPI (4′,6-diamidino-2-phenylindole) stained nuclei appear greenish-blue (variation in hue).

Correcting both inter- and intra-image variation is critical for both the segmentation and classification of ROIs. Regions of a particular structure class may be under- or over-segmented due to their location within the FOV of an image and the image to which they belong. The number of pixels in a region due to the quality of segmentation, coupled with the differences in hue, saturation, and value of pixels within a region can greatly affect the feature metric calculations. The feature metric variance for regions within a class can, in turn, then affect the performance of the classification results. Figure 2 shows the effect of these preprocessing steps on homogenizing images in an image set.

To correct the intra-image variation due to the single light source, we create a mask from all the blue pixels in the image. Blue pixels represent the DAPI-stained cell nuclei and are the most common and evenly distributed color in the microscopy images we analyzed. The blue mask is applied to the value channel of the image and the masked value data is fitted using a bivariate Gaussian. The Gaussian fit is then inverted and sampled at each pixel coordinate of the image, and then summed with the original unmasked value channel. This process is repeated for every image in the image set.

Following value gradient correction, the inter-image color variation is corrected, starting with the selection of a reference image from the image set. The blue mask is extracted from each gradient corrected image to calculate the mean hue value of those pixels. The image with a mean hue closest to the “true blue” of 120 (on a 180-value hue scale) is chosen as the reference image. The images are converted to International Commission on Illumination (CIE) L*a*b space to transfer the color profile from the reference image to the other images in the image set^35^.

### Generation of the signal mask

The homogenized images are processed to enhance visual contrast of the structures targeted by the probes. In the LungMAP images, blue indicates DAPI staining of nucleated cells and is not utilized as a fluorescent probe color. By clipping the blue channel in RGB color-space to the maximum of the red and green channels, areas of high blue intensity are suppressed. This reduces the saturation and value of regions not targeted by any fluorescent probes, thus increasing visual contrast of the borders in adjacent anatomical structures.

Each contrast enhanced image is then converted to a hue, saturation, and value (HSV) color space, where the value channel is used as input to a difference of Gaussians routine to generate a binary signal mask that highlights boundaries in visually contrasting areas. The boundaries of neighboring structures within the signal mask are often connected, and single structures may be incomplete, preventing the use of the signal mask to segment regions directly. Instead, the HSV image is used for segmentation of candidate regions of interest (ROI) based on color and saturation, and those regions are dilated and compared to the signal mask to infer the boundaries of the final ROIs (Figure 3).

### Customizable segmentation pipeline

The segmentation pipeline uses a customizable sequence of segmentation stages. Regions generated in each stage are removed from subsequent stages, i.e. the residual is used as the input for the next stage ensuring that there are no overlapping regions generated (Figure 4). There are two types of segmentation stages, one based on color and the other based on saturation. The type and number of segmentation stages is configurable in the micap Python library by a list of dictionaries, where each dictionary defines a segmentation stage, and each stage is executed in the order it appears in the list. Configurable parameters for each stage are used to target structures of a particular size range, and include a 2-D blur kernel size, a minimum ROI size, and a maximum ROI size. Color stages require an additional parameter, listing the color names to use for targeting specific structures containing a combination of fluorescent probes in addition to the targeted size range. The colors listed in a single-color stage correspond to the color definitions described previously, and their HSV ranges are combined to create masks targeting structures using that particular combination of probe colors.

### Post-processing of segmented regions

The contours found at each segmentation stage are accepted or rejected by iteratively dilating them against the signal mask. This is necessary as the individual segmentation stages target probes, which are generally bound to sub-cellular components, and the resulting regions may be under-segmented with respect to the complete cellular and structural boundaries. As described previously, the signal mask provides more accurate boundaries for regions of visual contrast but cannot be used to directly generate regions of interest due to its inter-connection with adjacent structures and broken borders of single structures. The final size for the ROI is determined by calculating the percentage of “on” pixels from the signal mask and choosing the iteration where the signal percentage falls off rapidly (Figure 3).

The contours generated by the automated segmentation stages often contain more detail than is necessary for quantitative analysis. This is manifested by an excessive number of vertices defining the perimeter of the region, resulting in a significant increase in the storage size of the arrays defining the regions and a less appealing visual appearance. Thus, the region boundaries are smoothed using the Douglas-Peucker algorithm^36^ as implemented in OpenCV^37^ to reduce the number of vertices, reducing storage volumes and increasing visual appeal while having an insignificant effect on the generated feature metrics passed on to the classification routine.

### Region classification

The pixel data in the post-processed regions is extracted and utilized to calculate the feature metrics described previously. The feature matrix is then scaled and passed to a classical machine learning pipeline for classification. After evaluating multiple classification strategies, including several deep learning neural networks, as well as more traditional machine learning techniques such as SVM, the most performant classifier was XG-Boost (XGB)^23^.

The classification pipeline returns the predicted label along with the probabilities for each structure class. The probabilities can then be used to optionally accept or reject regions, for example, by choosing the class with maximum posterior probability. As noted above, if any of the images in the training data were comprehensively segmented, the prediction results will also include the probability of belonging to the image ‘background’. Since all classified regions will likely have a category where the probability is highest (probability ties are rare), the inclusion of a background class reduces the occurrence of false positives due to partial segmentation or gross over-segmentation of structures, as well as ROIs that were generated in regions where no structure exists. Regions classified as “background” are then easily filtered from the final results.

### Cell segmentation

Once regions are classified, an optional final step in the pipeline is to sub-segment the structure into its component cells. Here, the ontology also plays a role in determining how to segment the structure into cells. The ontology provides the list of tissues and cell types found within a structure as well as whether the probes present in the structure are on the periphery or in the interior of the structure. Together with the intracellular DAPI staining and the fact that black represents non-cellular areas, the non-peripheral probe colors are used to segment areas most likely to contain cells. We developed a recursive binary segmentation procedure to identify nested structures within a region, where each region is recursively partitioned into two sections using spectral clustering, stopping at a configurable minimum size (Figure 4b). The cell contours are then returned from this recursive binary spectral clustering for display or further analysis.

## Acknowledgements

We thank the University of Rochester LungMAP Human Tissue Core (U01HL122700), Joe Kitzmiller and Jeff Whitsett for providing the original images from Cincinnati Children’s Hospital Medical Center (https://research.cchmc.org/lungimage/), Gail Deutsch and Jerry Kirchner for discussion and suggestions, Jeramie Williford and Framers’ Workroom of Jenkintown, PA for their professional printing services. Special acknowledgment goes to the families of donors for their generous and irreplaceable contributions to the successes of the LungMAP program. Funding for this study was provided by the National Heart, Lung & Blood Institute grant 5U01-HL122638 for “Molecular Atlas of Lung Development – Data Coordinating Center”.

## Authors’ contributions

AMM developed the application ontology. SW provided testing and queries for the ontology and was the primary developer of the software libraries created for this project. BN researched and implemented current state of the art deep learning architectures as well as provided statistical support in reporting the results. MAP and NG provided support in generating SPARQL queries and accessing the BREATH database. CBH contributed to the LungMAP ontology development, development of the image annotation tool - helping guide image annotation and facilitated meeting and discussions during the development of the automated image analysis system. RSM annotated images and assisted discussions regarding lung structure. BA provided framework considerations and initial models of lineage and sub-lineage temporal and morphogenesis trajectories within which models of ontology-mapped cell types, developmental cell-cell transitions, and intercellular adjacency relationship structures for segmentation, classification, and featurization of multi-marker developing lung molecular histology data could be analyzed. SEW is the histopathology expert of the developmental lung who provided the “ground truth” for the IF images tested. LY implemented and extended algorithms for image segmentation in the early stage of software development. SP provided domain expertise and steered the overall direction of the project. CC conceived of the methodological approach, directed the work and prepared the first draft of the manuscript. All authors edited and approved the manuscript.

## Competing interests

The authors declare that they have no competing interests.

**Supplemental Table 1:**
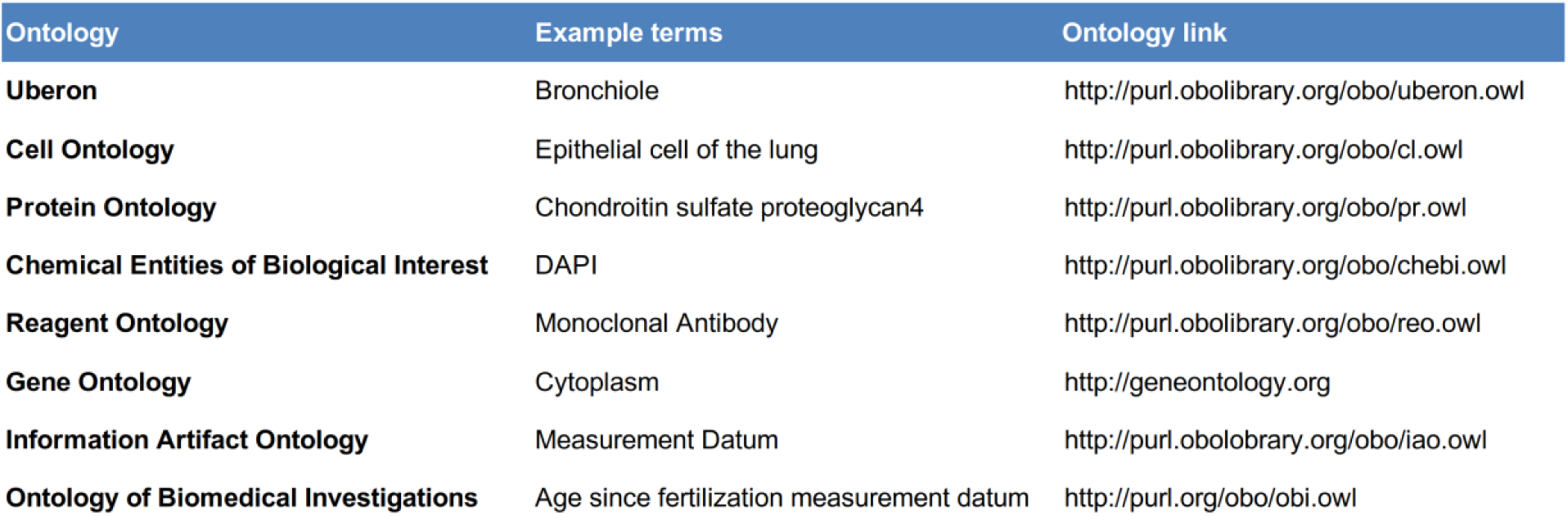
Examples of terms from foundational ontologies found in the application ontology.

**Supplemental Table 2:**
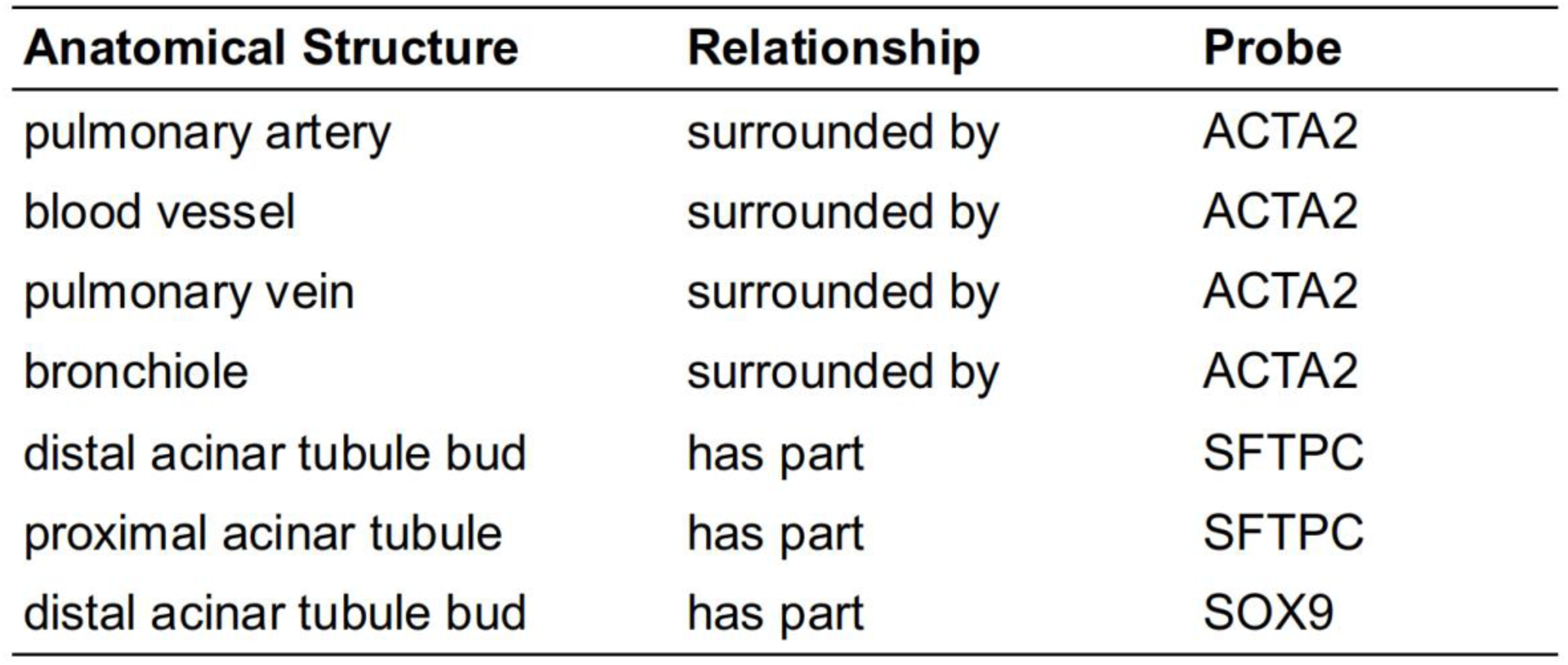
Ontology query results for the probes present in an image, demonstrating the linkage between probes and anatomical structures, as well as their contextual information regarding location where they are present in the structure.

**Supplemental Table 3:**
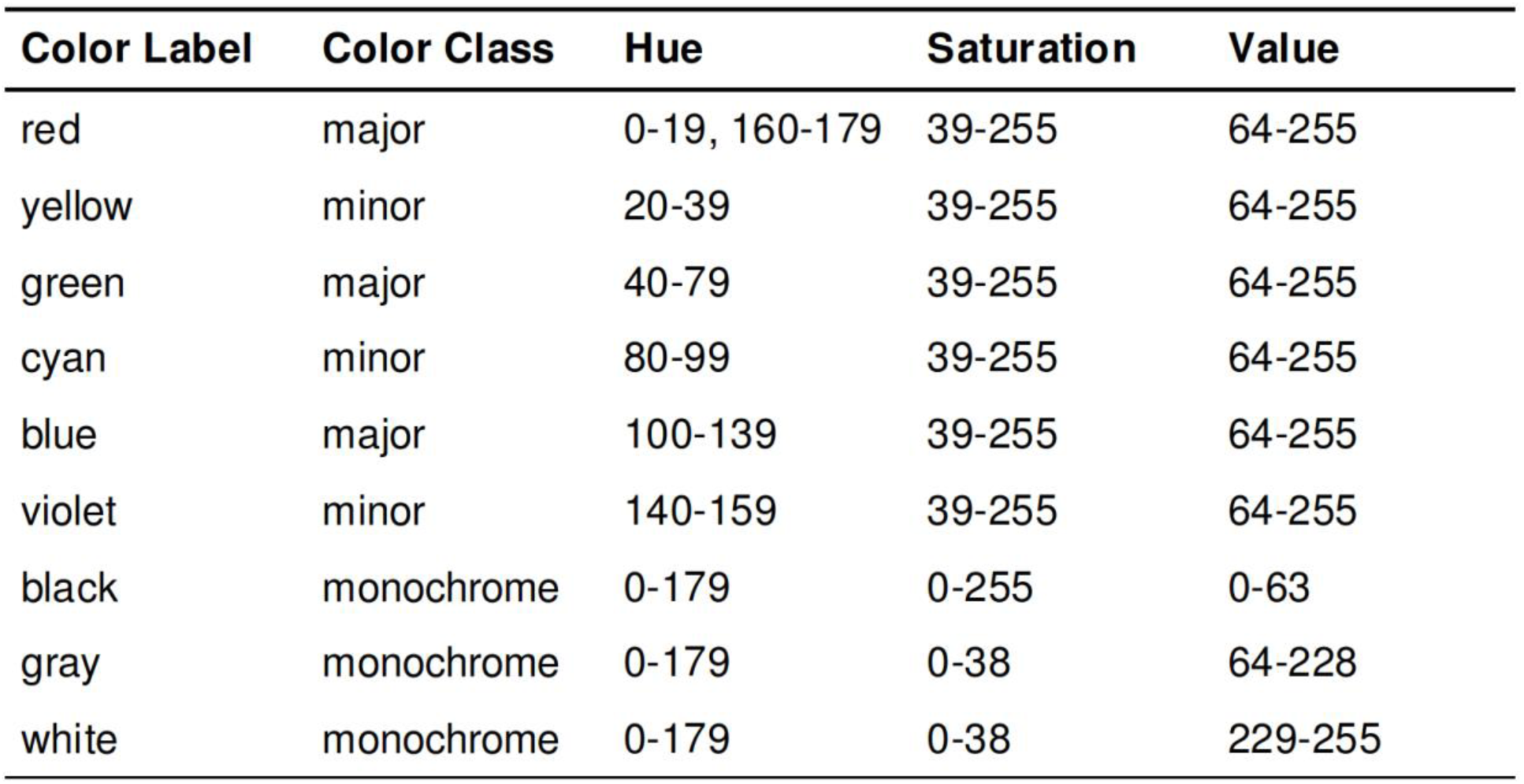
Color definitions utilized in the image analysis pipeline. Colors are defined by partitioning the complete HSV color space such that each HSV value occurs in only a single color label. The hue range spans 180 values, saturation spans 256 values, and value (intensity) spans 256 values. Color labels are classified into 3 groups: major colors spanning 40 hue values, minor colors spanning 20 hue values, and monochrome colors.

