## Supplemental Table 1 for "Ontology-guided segmentation and object identification for developmental mouse lung immunofluorescent images"

Supplemental Table *1*: Examples of terms from foundational ontologies found in the application ontology

| Ontology | Example terms | Ontology link |
|---|---|---|
| **Uberon** | Bronchiole | http://purl.obolibrary.org/obo/uberon.owl |
| **Cell Ontology** | Epithelial cell of the lung | http://purl.obolibrary.org/obo/cl.owl |
| **Protein Ontology** | Chondroitin sulfate proteoglycan4 | http://purl.obolibrary.org/obo/pr.owl |
| **Chemical Entities of Biological Interest** | DAPI | http://purl.obolibrary.org/obo/chebi.owl |
| **Reagent Ontology** | Monoclonal Antibody | http://purl.obolibrary.org/obo/reo.owl |
| **Gene Ontology** | Cytoplasm | http://geneontology.org |
| **Information Artifact Ontology** | Measurement Datum | http://purl.obolobrary.org/obo/iao.owl |
| **Ontology of Biomedical Investigations** | Age since fertilization measurement datum | http://purl.org/obo/obi.owl |
