## Supplemental Table 2 for "Ontology-guided segmentation and object identification for developmental mouse lung immunofluorescent images"

Supplemental Table 2: Ontology query results for the probes present in an image, demonstrating the linkage between probes and anatomical structures, as well as their contextual information regarding location where they are present in the structure.

| Anatomical Structure | Relationship | Probe |
|---|---|---|
| pulmonary artery | surrounded by | ACTA2 |
| blood vessel | surrounded by | ACTA2 |
| pulmonary vein | surrounded by | ACTA2 |
| bronchiole | surrounded by | ACTA2 |
| distal acinar tubule bud | has part | SFTPC |
| proximal acinar tubule | has part | SFTPC |
| distal acinar tubule bud | has part | SOX9 |
