## Supplemental Table 3 for "Ontology-guided segmentation and object identification for developmental mouse lung immunofluorescent images"

Supplemental Table 3: Color definitions utilized in the image analysis pipeline. Colors are defined by partitioning the complete HSV color space such that each HSV value occurs in only a single color label. The hue range spans 180 values, saturation spans 256 values, and value (intensity) spans 256 values. Color labels are classified into 3 groups: major colors spanning 40 hue values, minor colors spanning 20 hue values, and monochrome colors.

| Color Label | Color Class | Hue | Saturation | Value |
|---|---|---|---|---|
| red | major | 0-19, 160-179 | 39-255 | 64-255 |
| yellow | minor | 20-39 | 39-255 | 64-255 |
| green | major | 40-79 | 39-255 | 64-255 |
| cyan | minor | 80-99 | 39-255 | 64-255 |
| blue | major | 100-139 | 39-255 | 64-255 |
| violet | minor | 140-159 | 39-255 | 64-255 |
| black | monochrome | 0-179 | 0-255 | 0-63 |
| gray | monochrome | 0-179 | 0-38 | 64-228 |
| white | monochrome | 0-179 | 0-38 | 229-255 |
